# Sex differences in task engagement and lapse rate during reward learning

**DOI:** 10.1101/2025.10.29.685451

**Authors:** C.G. Aguirre, J.H. Woo, L. Alhabbal, T. Fujioka, R. Moore, T. Ye, J.J. Castrellon, A. Soltani, A. Izquierdo

**Author notes:** **Correspondence**: Alicia Izquierdo, Ph.D. equal contribution, co-first authors (C.G. Aguirre, J.H. Woo). **Author contributions:** CGA and AI designed the research; CGA, LA, TF, TY, and RM conducted the research; CGA, JHW, AS and AI analyzed the data; CGA, JHW, JJC, AS, and AI interpreted the data; AS and AI acquired funding for the project; CGA, JHW, and AI wrote the paper; all authors edited the final version.

## Abstract

Our understanding of sex differences in reward learning has been limited due to the predominant study of males. Here, we evaluated sex differences in flexible learning in two domains: the learning of stimulus-and action-based associations and their reversals. During action-based learning, rats selected between two identical visual stimuli presented on a touchscreen, where the spatial location predicted a higher probability of reward. For stimulus-based learning, rats chose between two distinct visual stimuli presented in pseudorandom spatial locations, one of which was associated with a higher probability of reward. Reversal phases involved switching reward contingency between the two actions or stimuli. We found that females did not differ in their discrimination or reversal learning accuracy compared to males, yet they collected fewer rewards and were more likely to omit trials than males in both domains. To gain a detailed understanding of differences across conditions, we modeled animals’ trial-by-trial choices using reinforcement learning (RL) models and examined their steady-state behavior to capture transitions between distinct behavioral states. Although the estimated parameters of the best-fitting RL model revealed some sex differences, the model that incorporated transitions between different behavioral states provided a better overall fit to the data. This model also revealed that across all reversal phases, females exhibited a higher transition-specific lapse rate than males, indicating greater task disengagement once there was no need for further learning. Together, our fine-grained analysis of behavior adds to a growing literature on sex differences in flexible reward learning.

## Introduction

There are persistent gaps in our understanding of sex differences in the context of reward-based learning and decision-making due to the sparsity of studies that include female animals. The emergence of sex-dependent effects in the few studies that have included both sexes has put into question the generalizability and interpretations of previous findings that only included males. There have been reports of sex differences in other measures that contribute to the decision-making process including patterns of exploration, use of adaptive strategies (1, 2), impulsivity (3), and sensitivity to negative feedback (4, 5). Importantly, studies that have included both sexes have not uncovered sex differences in learning accuracy in either action(spatial)-based (2) or stimulus-based (1, 4) learning tasks.

A common way to study flexible reward learning and decision-making is through reversal learning, which measures a subjects’ ability to form flexible associations between stimuli and/or actions with their outcomes (6–8). Importantly, reversal learning paradigms can be used to probe learning under uncertain conditions in a variety of modalities (e.g. visual, spatial, olfactory). Additionally, when fine-grained trial-by-trial data is collected, such behavioral methods allow us to gather information about attention, deliberation, and motivation through initiation, choice, and reward collection latencies, respectively, and to compute sensitivity to positive and negative feedback through Win-stay and Lose-shift metrics (9, 10). These performance measures may increase our sensitivity to detect sex differences by allowing us to capture dynamic behavioral changes within a session and across time.

Sex differences have been described in several components of learning, such as the employment of adaptive strategies, with female mice displaying more Win-stay behaviors than males on a spatial-based two-armed bandit probabilistic task (2). Some groups have also uncovered that males and females differ in explore/exploit behaviors, with female mice exploring less overall, but exploiting the better option much earlier in learning compared to males. Conversely, males tend to explore more overall and exploit the better option later in learning than females (1, 2). Additionally, females tend to be more sensitive to negative feedback(4, 5, 11, 12), exhibiting longer trial initiation latencies following unrewarded outcomes, in response to negative feedback (4). Altogether, these findings demonstrate that males and females may differ in their employment of adaptive strategies, as well as sensitivity to positive and negative feedback, which can influence learning.

Here, we evaluated sex differences in the context of flexible learning using stimulus-and action-based reversal learning tasks. Animals were required to learn to associate either a spatial side (i.e. right or left) or a visual stimulus (i.e. fan or marble) with reward or no reward. After meeting criterion, the reward contingencies were reversed, such that the previously rewarded side or stimulus was no longer rewarded, and vice versa. Additionally, we varied the level uncertainty in the environment by testing them on both deterministic (100/0) and probabilistic (90/10) reward outcomes. Overall learning and accuracy were assessed across trials (for action learning) and sessions (for stimulus learning), along with performance measures such as use of adaptive strategies (i.e., win-stay, lose-shift), omissions, and latencies, to better assess potential sex differences. To capture the mechanism underlying sex differences in action-based reversal learning, we utilized computational models based on reinforcement learning (RL) and also adapted a computational method based on sigmoidal transition curves (13) to characterize behavioral transitions, and modified it to include distinct lapse rates, both prior to and following the transition in reversal. Overall, the combination of these methods enabled a more detailed assessment of sex differences in the post-reversal dynamics of learning.

## Materials and Methods

Timeline of experimental procedures is presented in **Figure 1A**.

**Figure 1.**
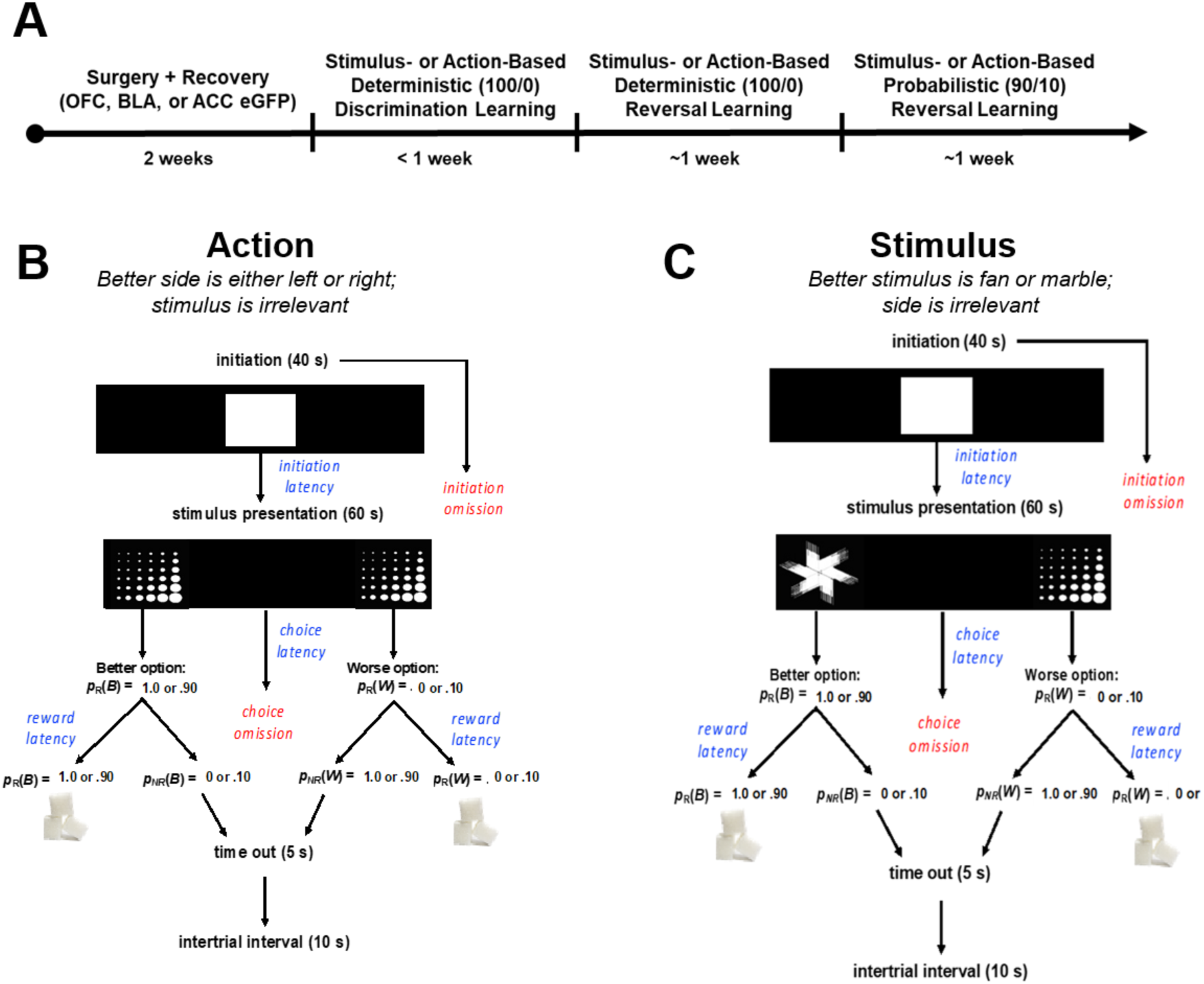
**Experimental timeline and reversal learning tasks.** (**A)** *Experimental Timeline.* Only animals with enhanced green fluorescent protein (eGFP) null virus in orbitofrontal cortex (OFC), basolateral amygdala (BLA), or anterior cingulate cortex (ACC) were included in these analyses. (**B**) *Action-based reversal learning task structure.* Animals initiated trials by nosepoking the white center square, after which two identical visual stimuli (fan or marble) were presented on the left and right side of the screen and whose spatial side was associated with either a 100/0 (deterministic) or 90/10 (probabilistic) reward probability. (**C**) *Stimulus-based reversal learning task structure.* Animals initiated trials by nosepoking the white center square, after which two different visual stimuli (fan and marble) were presented on the left or right side of the screen, with each stimulus associated with either a 100/0 (deterministic) or 90/10 (probabilistic) reward probability.

### Subjects

Animals were adult (N=32, 15 females) Long-Evans rats (Charles River Laboratories) average age post-natal-day (PND) 65-85 at the start of experiments, with a 240g body weight minimum for females and 280g body weight minimum for males at the time of surgery. Subjects for these experiments were male and female surgerized null virus control animals from Aguirre et al. (14)(n=14) and additional cohorts of animals of the same age (n=18) (see details on viral surgeries below). All rats underwent a 3-day acclimation period immediately upon arrival, in which they were pair-housed and given food and water *ad libitum*, with no experimenter interference. After the 3-day acclimation period, animals were handled over 5 consecutive days for 10 min each, and continued to be provided food and water *ad libitum*. Following the handling period, animals were individually-housed in standard vivarium housing conditions (room temperature 22–24° C) with a reverse 12 h light/dark cycle (lights off at 6am). Animals underwent surgery and then were first tested on pretraining schedules after a week of post-operative care. Following pre-training, animals were tested on discrimination learning followed by reversal learning, at which point they were at the minimum 3-week expression time for the null virus and could be administered clozapine-N-oxide (CNO) or saline as vehicle (VEH). To include all animal data, we entered drug as a covariate in order to account for any potential behavioral effects following CNO administration. All procedures were conducted in accordance to the recommendations in the Guide for the Care and Use of Laboratory Animals of the National Institutes of Health and with the approval of the Chancellor’s Animal Research Committee at the University of California, Los Angeles.

### Surgery

#### Viral Constructs

Rats were singly-housed and remained in their home cages for 3-4 weeks prior to testing. As part of a viral control group, they were prepared with an enhanced Green Fluorescent Protein (eGFP) on a CaMKIIa promoter (AAV8-CaMKIIa-eGFP, Addgene #176015), infused bilaterally into regions of the frontal cortex or amygdala. The eGFP vector was injected in BLA neurons (n=8) [(AP =-2.5; ML= ±5; DV = −7.8 (Vol. 0.1 µl),-8.1 (Vol. 0.2 µl)], vlOFC neurons (n=6) [AP = +3.7; ML= ±2.5; DV = −4.6, (Vol. 0.2 µl); AP= 4; ML= ±2.5; DV= −4.4, (Vol. 0.15 µl)], or ACC neurons (n=18) [AP = +3.7; ML= ±0.8; DV =-2.6 (Vol. 0.3 µl)], measured from bregma and infused at a rate of 0.1 µl/min.

#### Surgical Procedure

Infusions of eGFP control virus were performed under isoflurane gas (1-5% in O_2_) anesthesia using aseptic stereotaxic techniques prior to any behavioral testing. During surgery, all animals were administered 5 mg/kg s.c. carprofen (NADA #141–199, Pfizer, Inc., Drug Labeler Code: 000069) and 1cc saline. After being placed in the stereotaxic apparatus (David Kopf; model 306041), an incision was made on the scalp and the scalp was retracted. The skull was leveled with a +/-0.3 mm tolerance on the anterior-posterior axis in order to ensure that bregma and lambda were in the same horizontal plane. Small holes were drilled in the skull above the infusion target. Virus was bilaterally infused at a rate of 0.1 µl per minute in target regions (see coordinates above), after which, 5 min were allowed to elapse before withdrawing the needle containing the virus.

### Histology

After completing the experiment, rats were euthanized with an overdose of Euthasol (Euthasol, 0.8 mL, 390 mg/mL pentobarbital, 50 mg/mL phenytoin; Virbac, Fort Worth, TX), were transcardially perfused, and their brains removed for histological processing. Brains were fixed in 10% buffered formalin acetate for 24 h followed by 5 days in a 30% sucrose solution. Afterward, the brains were sectioned into 40-µm coronal sections, mounted onto slides, and cover-slipped with DAPI. To visualize eGFP expression in cell bodies, slices were visualized using a BZ-X710 microscope (Keyence, Itasca, IL), and analyzed with BZ-X Viewer and analysis software.

### Food Restriction

Rats were placed on food-restriction five days prior to any behavioral testing, with males given 12-14 grams of chow a day, and females given 10-12 grams/ day. Food restriction level was maintained throughout behavioral testing. Water remained freely available in the home cage. Animals were weighed at least every other day, and their weight was monitored so as to not fall below 85% of their maximum, free-feeding weight.

### Drug administration

Thirty minutes prior to behavioral testing, systemic administration of clozapine-N-oxide, CNO (i.p., 3mg/kg in 95% saline, 5% DMSO) or saline vehicle (VEH) was injected intraperitoneally. We followed a within-subject design for reversal learning, such that all rats received CNO and VEH injections in counterbalanced order. Thus, if a rat received VEH on the first reversal (R1), it was administered CNO on the second reversal (R2), with the same drug order for the third reversal (R3) and the fourth reversal (R4), or vice versa, as reported in Aguirre et al. 2024(14).

### Behavioral Testing

*Pretraining.* Behavioral testing was conducted in operant conditioning chambers containing an LCD touchscreen and a sucrose pellet dispenser on the opposing side. All chamber equipment was operated using customized ABET II TOUCH software (Lafayette Instrument Co., Lafayette, IN). The pretraining protocol was adapted from established procedures(15), consisting of a series of training schedules: Habituation, Initiation Touch to Center (ITC), and Immediate Reward (IM). These pretraining schedules were designed to train rats to collect sucrose pellet rewards from the magazine, nose poke a center stimulus to initiate a trial, and to select a stimulus located on either the left or right side of the screen to obtain a reward. Pretraining schedules and criterion have been reported in more detail elsewhere(14). After completing all pretraining schedules, rats were advanced to the discrimination (initial) phase of either the action-and/or stimulus-based reversal learning task, with the task order counterbalanced (**Fig. 1**).

*Action-based deterministic discrimination learning.* After completion of all pretraining schedules, rats were advanced to the discrimination (initial) phase of the action-based task (**Fig. 1B**). Rats were required to touch a white square stimulus on the center of the screen (40 seconds) to initiate a trial, after which they were presented with the same two visual stimuli (i.e., marble or fan) on the left and right side of the screen (60 seconds). Rats could nosepoke either the spatial side that was rewarded with a sucrose pellet, or the spatial side that was unrewarded, followed by a 10 s inter-trial interval (ITI). If the trial was unrewarded, a time-out of 5 s occurred prior to the ITI. Rats had to choose the correct side 75% of the trials or more, collect at least 60 rewards during a 60 min testing session for two consecutive days, to meet criterion. Animals were not administered any CNO or VEH injections during discrimination learning. After meeting criterion, rats were advanced to the first reversal phase in the next testing session.

*Action-based reversal learning.* After the discrimination learning phase, the rats underwent four reversals. Rats were injected intraperitoneally with either 3 mg/kg of CNO or VEH 30 min before each reversal testing session. For the first two deterministic reversals, the side previously associated with the higher reward probability (p_R_(B)=1.0) was now associated with a lower reward probability (p_R_(W)=0.0), and vice versa. The criterion was the same as the deterministic discrimination phase. After reaching the criterion for the first deterministic reversal phase (i.e., R1), the rats advanced to the second deterministic reversal phase (i.e., R2) beginning on the next testing session. Rats that had previously received VEH during the first reversal would now receive CNO injections during the second reversal, and vice versa. After completing two deterministic reversals, rats underwent two more reversals under probabilistic conditions (reversals 3 and 4), where the side with the highest reward probability was associated with p_R_(B)=0.9, and the side with the lower reward probability was associated with p_R_(W)=0.1.

*Stimulus-based deterministic discrimination learning.* After completion of all pretraining schedules, rats were advanced to the discrimination (initial) phase of learning of the stimulus-based task (**Fig. 1C**). Rats were required to touch a white square stimulus on the center of the screen to initiate a trial, after which they were presented with two different visual stimuli (i.e., marble and fan). Stimuli were randomly assigned as the correct or incorrect stimulus, associated with either a p_R_(B)=1.0 or p_R_(W)=0.0 probability of reward, respectively. Rats would nosepoke the stimulus of their choosing and depending on its reward probability, received a sucrose pellet reward or 5 s time-out period, followed by a 10 s ITI. The criterion was the same as the action-based task. Rats were given a maximum of 10 days to achieve criterion and then were advanced to the first reversal learning phase. Animals were not administered either CNO or VEH injections during discrimination learning.

*Stimulus-based reversal learning.* After the discrimination learning phase, the rats underwent four reversals. Rats were injected intraperitoneally with either 3 mg/kg of CNO or VEH 30 min before each reversal testing session. The stimulus previously associated with the higher reward probability (p_R_(B)=1.0), was now associated with a lower reward probability (p_R_(W)=0.0), and vice versa. The criterion was the same as the deterministic discrimination phase. After reaching the criterion for the first deterministic reversal phase (i.e., R1), the rats advanced to the second deterministic reversal phase (i.e., R2) beginning on the next testing session. Rats that had previously received VEH during the first reversal would now receive CNO injections during the second reversal, and vice versa. After completing two deterministic reversals, rats underwent two more reversals under probabilistic conditions (reversals 3 and 4), where the stimulus with the highest reward probability was associated with p_R_(B)=0.9, and the side with the lower reward probability was associated with p_R_(W)=0.1.

## Data Analyses

MATLAB (MathWorks, Natick, Massachusetts; Version R2022b) was used for all statistical analyses and figure preparation. For each Action and Stimulus-based learning dataset, we fitted Generalized Linear Models (GLMs; *fitglme*) to examine the effects of sex on the probability of choosing the better option across trials or sessions. For selection of the best-fitting model, we used the GLM formula with the lowest Bayesian Information Criterion (BIC). Mixed-effects GLMs were conducted for each task separately, for each outcome variable, and with trials and sessions as within-subject factors, sex as a between-subject factor, and individual rat as a random factor, along with other within-subject factors such as reversal number (1–4) and drug (i.e. VEH, CNO) as covariates. Importantly, CNO was not expected to have an effect on learning or behavior since all subjects included here were null virus (eGFP) controls. For learning accuracy, session and trial were included in the models as a moderator that could interact with sex using one the following formulas: (Action-based) γ∼[1+sex*session*trial+reversal+drug+(1+session*trial+reversal|rat)]; (Stimulus-based) γ∼[1+sex*session*trial+reversal+drug+(1+session*trial+reversal|rat)].

For performance measures such as latencies, rewards collected, and initiation omissions, session was not included in these formulas as measures were either summed or averaged across sessions. Thus, the formulas were as follows for both Action-and Stimulus-based tasks: γ∼[1+sex+reversal+drug+(1+reversal|rat)]. Since learning reached asymptote after 5-days for stimulus-based reversal learning, only the first 5 testing sessions were included in the GLM for learning in that domain. For action-based learning, only trials included in the first two testing sessions were included in the GLM analyses, since most rats only required two days to meet criterion. Significant reversal number and/or sex interactions were further analyzed with a narrower set of fixed factors and Bonferroni-corrected post-hoc comparisons.

Dependent measures for learning included probability of choosing the correct or better option, number of rewards collected, median initiation latencies and number of omissions (latency to initiate a trial and failure to initiate a trial, respectively), median correct and median incorrect choice latencies (latency to select the correct or better stimulus or spatial side and latency to select the incorrect or worse stimulus or spatial side, respectively), median reward latencies (latency to collect the reward), probability of Win-stay, and probability of Lose-shift. Probability of Win-stay and Lose-shift adaptive strategies were calculated such that each trial was classified as a *win*, when the animal received a sucrose pellet and as a *loss* if there was no reward delivered. Statistical significance was noted when p-values were less than 0.05. All Bonferroni post-hoc tests were corrected for number of comparisons. Outliers greater than two standard deviations above the mean were removed from the dataset.

### Analysis of choice behavior using reinforcement learning (RL) models

To capture the difference in learning and choice behavior between male and female rats, we utilized two simple conventional RL models as investigated in our previous study (14). The subjective estimate of reward for each choice option (Q*_i_*) was updated on a trial-by-trial basis based on the discrepancy between actual and expected reward values. In the first model, referred to as *RL*, the value estimate of the chosen option (*Q_C_*) was updated as follows:

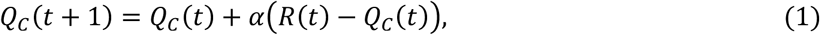

where *R*(t) indicates reward outcome on trial *t* (1 if rewarded, 0 otherwise), and 𝛼 is the learning rate controlling the amount of update. The second model, referred to as *RL_decay_,* additionally updated the value of the unchosen option (*Q_U_*) as follows (16):

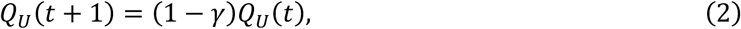

where 𝛾 is the decay rate controlling the amount of passive decay in the value of the unchosen option. For this model, on omitted trials where no choice was made, both left and right options were considered unchosen, and their value estimates decayed passively following this equation.

Both models used the following soft-max function to compute the probability of choosing a leftward option (*P_L_*):

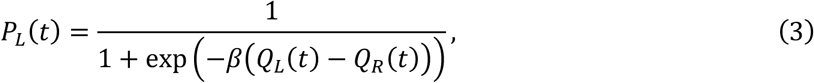

where *Q_L_* and *Q_R_* indicate the value estimates of left and right option, respectively, and 𝛽 is the inverse temperature or choice sensitivity governing the extent to which higher-valued options are consistently selected. The likelihood for each session was given by:

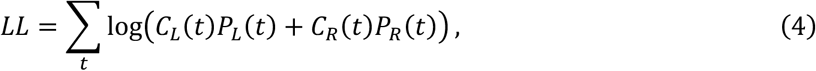

where *C_L_* and *C_R_* indicate whether the animal has chosen left or right option, respectively (1 if chosen, 0 otherwise), and *P_L_* and *P_R_* indicate choice probability obtained from Eq. (3) on trial *t* (𝑃_R_ = 1 – 𝑃*_L_*). To assess the goodness-of-fit, we computed the Akaike information criterion (AIC) for each session as follows:

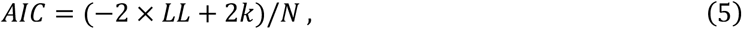

where *k* is the number of free parameters in the model (two for *RL* and three for *RL_decay_*), and *LL* is the best (i.e., maximum) log-likelihood value selected from the iteration for the given session as computed in Eq. (4). *N* indicates the number of trials in the given session. Since each session differed in the number of total trials, we normalized the AIC value by the number of trials. We used the standard maximum likelihood estimation method (using *fmincon* in MATLAB) to fit choice data and estimate the parameters for each session of the experiment. For each run, we repeated 100 different initial conditions selected from evenly spaced search space to avoid local minima.

### Capturing behavioral state transitions

In addition to the RL models above, we also examined behavioral transitions following reversals using a model based on a sigmoidal transition curve (13). Specifically, we considered two sigmoidal transition functions that involved the latency, the speed, and the lapse rate of transition after reversal in each session. The first sigmoidal curve, referred to as *SC1*, described the probability of choosing the better rewarding option (*P_Better_*) as follows:

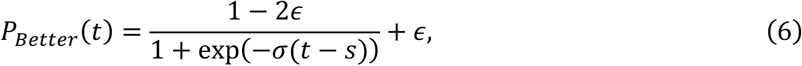

where 𝜎 is the slope of the sigmoidal curve controlling the steepness (speed) of transition, *s* is the offset of the curve specifying the latency of transition, and 𝜖 is the lapse rate which specifies the rate of error after the behavioral transition.

Because the performance could potentially differ before and after the choice transition, we also tested the second sigmoidal curve, referred to as SC2, to dissociate the lapse rates before and after the transition has occurred (as illustrated in **Fig. 4A**). This two-lapse function was written as:

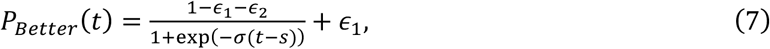

where 𝜖_1_ and 𝜖_2_ represent the lapse rates before and after the choice transition, respectively. Note that each lapse rate indicates the rate at which the animal’s choice deviates from the options assumed to be subjectively dominant during a given period, i.e., the worse option before transition and the better option after transition.

As was done for the RL models, we fitted the sigmoidal curves to the choice data and estimated the parameters for each session of the experiment. To obtain the goodness-of-fit comparable with that of the RL models, we defined the log-likelihood with each session as:

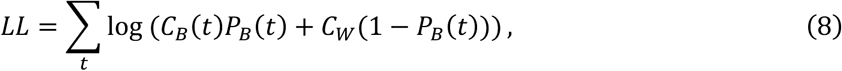

where *C_B_* and *C_W_* indicate whether the animal has chosen better or worse option, respectively (1 if chosen, if 0 otherwise) on trial *t*. Omitted trials were excluded from the fit. *P_B_*(*t*) is the choice probability for better option as calculated above in Eq. (6) or (7). Note that this definition is equivalent to the log-likelihood obtained from the RL model in Eq. (4). To quantify the goodness-of-fit to trial-by-trial choice data, we computed the AIC normalized by number of trials per session, for sigmoid functions according to Eq. (5) (*k* = 3 for *SC1*, and *k* = 4 for *SC2*).

For probing sex differences on model-estimated parameters and model-dependent measures (e.g., performance after behavioral transitions), we used a mixed-effects model with sex as the main predictor and subject as a random factor, with the following formula: γ ∼ [1+sex + (1|rat)]. This mixed-effects GLM provided the best fit with the lowest BIC. For correlations between lapse rates and number of omissions, we used the estimated offset parameter (*s*) for each session and counted the number of omissions from trials only after this offset. Pearson correlation coefficients were computed separately for males and females.

## Results

### Females and males learn initial discriminations at comparable rates

To assess the speed of initial acquisition, we used GLM formulas to determine the major predictors of learning across both action and stimulus domains. Formulas were identical to those used for subsequent reversal learning except that reversal number and drug did not apply (i.e. animals were not injected in discrimination) and thus these were excluded in the model. Both males and females learned initial discriminations comparably across trials and sessions: There were no sex differences in discrimination learning accuracy (i.e., the probability of choosing the correct side) in the action-based task (β_sex_ =-0.14, *p* = 0.26; **Fig. 2A**) or in the stimulus-based task (i.e., probability of choosing the correct visual stimulus; β_sex_ =-0.05, *p* = 0.39; **Fig. 2C**). In action-based learning, there was a significant effect of session (β_sessions_ = 0.21, *p* = 4.56e-06) and trials (β_trials_ = 0.002, *p* = 4.29e-07) and a significant session*trial interaction (β_session*trial_ =-0.001, *p* = 5.61e-04). In stimulus-based learning, there was only a significant session*trial interaction (β_session*trial_ = 0.0003, *p* = 0.02). Rats exhibited much more difficulty learning the stimulus-based task regardless of sex with only 65% of rats meeting criterion for initial discrimination learning within 10 sessions. Nevertheless, there were marginal improvements by session for this type of learning (β_sessions_ = 0.03, *p* = 0.05).

**Figure 2.**
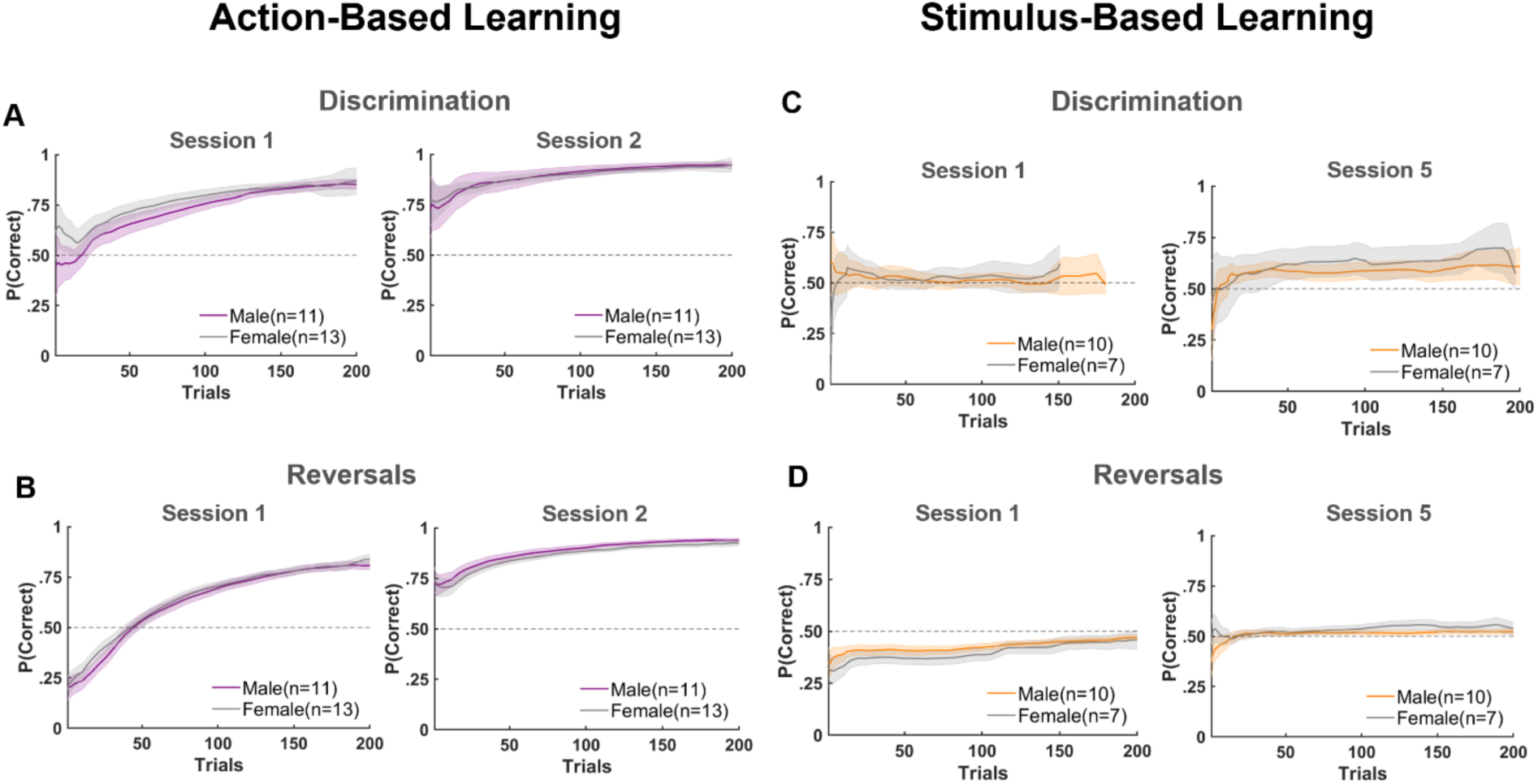
No sex differences in action-and stimulus-based discrimination or reversal learning. **(A)** There were no sex differences in action-based discrimination learning (i.e., the probability of choosing the correct side). Both males and females increased their accuracy across sessions and trials. **(B)** There were also no sex differences in action-based deterministic (100/0) or probabilistic (90/10) reversal learning. Both males and females increased their accuracy across sessions and trials. **(C)** There were no sex differences in stimulus-based discrimination (100/0) learning. **(D)** There were no sex differences in stimulus-based reversal learning. Solid lines depict group averages across all reversals and shading represents the Standard Error of the Mean (SEM).

As for performance measures, in action-based discriminations females omitted more trials (β_sex_ =-21.16, *p* = 0.001) and collected fewer rewards (β_sex_ = 78.85, *p* = 0.02) than males. There were however no sex differences in any of the latencies measured, including initiation latencies (*p* = 0.57), correct choice latencies (*p* = 0.52), incorrect choice latencies (*p* = 0.17), or reward latencies (*p* = 0.54). For stimulus-based discriminations, females took longer to initiate trials than males (β_sex_ =-1.22, *p* = 0.03), but there were no sex differences in the number of initiation omissions (β_sex_ = 6.10, *p* = 0.70) or rewards collected (β_sex_ = 38.81, *p* = 0.52). There were also no sex differences in any of the other latencies measured, including correct choice latencies (*p* = 0.70), incorrect choice latencies (*p* = 0.69), or reward latencies (*p* = 0.64).

### Females and males learn reversals at comparable rates

Both females and males learned action-based reversals comparably (β_sex_ =-0.04, *p* = 0.63), with both sexes increasing their accuracy across sessions (β_sessions_ = 0.39, *p* = 3.06e-22) and trials (β_trials_ = 0.01, *p* = 2.89e-31). There was also a significant session*trial interaction (β_session*trial_ =-0.002, *p* = 9.97e-14), indicating that learning of the new contingency following a reversal occurred mainly within the first session. On average animals achieved the 75% criterion for both session 1 (M±SEM: 0.81±0.08) and session 2 (0.93±0.10). Learning curves are shown collapsed across reversals (**Fig. 2B**).

Females and males learned stimulus-based reversals very slowly yet comparably (β_sex_ = 0.06, *p* = 0.23), with both sexes increasing their accuracy across sessions (β_sessions_ = 0.03, *p* = 0.001) and trials (β_trials_ = 0.001, *p* = 0.01). However, unlike action-based reversals, animals exhibited difficulty learning stimulus-based reversals with only 44% reaching a 75% criterion on deterministic reversals, and only 15% of rats learning probabilistic reversals within 10 sessions. Furthermore, we found that animals on average performed below chance levels for session 1 (M±SEM: 0.42±0.01) and only slightly above chance for later session 5 (0.53±0.07). Mean accuracy (i.e., probability of choosing the correct or better option) is shown for session 1 and session 5 collapsed across reversals (**Fig. 2D**).

### Males collect more rewards than females during reversal learning

We also assessed the influences of sex on successful reward procurement in the two different domains. Males collected more rewards than females in both action-based reversals (β_sex_ = 102.31, *p* = 4.75e-06) (**Fig. 3A**) and stimulus-based reversals (β_sex_ = 152.02, *p* = 5.59e-05) (**Fig. 3D**).

**Figure 3.**
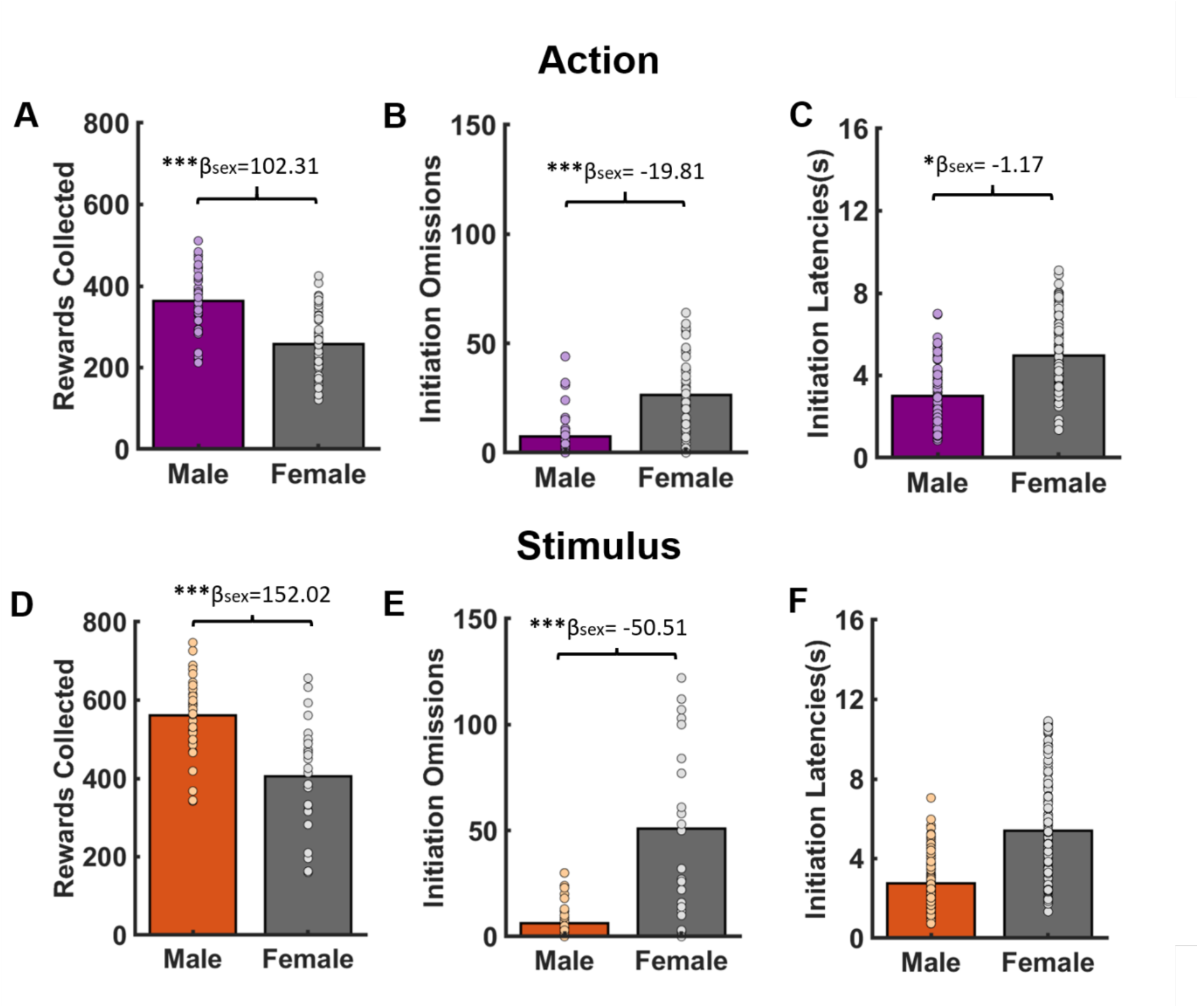
**Females collect fewer rewards and omit more trials than males in both action-and stimulus-based reversals**. **(A)** Males collected more rewards than females in action-based reversal learning. **(B)** Females omitted more trials than males in action-based reversal learning. **(C)** Females take longer to initiate trials than males in action-based reversal learning. **(D)** Males collected more rewards than females in stimulus-based reversal learning. **(E)** Females omitted more trials than males in stimulus-based reversal learning. **(F)** There were no sex differences in latencies to initiate trials in stimulus-based reversal learning. Asterisks indicate significant difference between females and males, *p<.01, **p<.001, ***p<.0001

**Figure 4.**
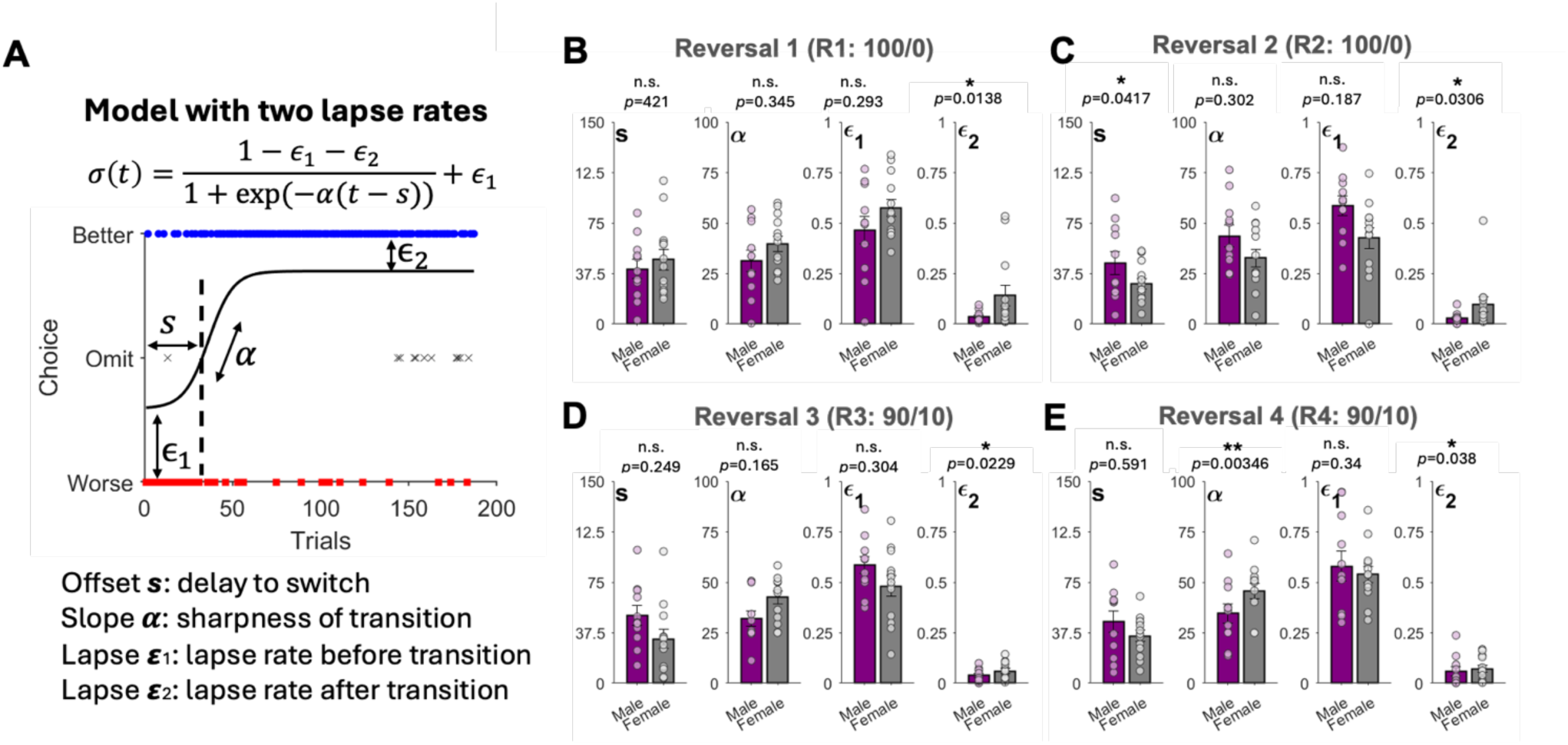
Females exhibit higher lapse rates after behavioral transition compared to males during action-based reversal learning. **(A)** Illustration of sigmoidal transition curve with two lapse rates. To characterize the animals’ switching dynamics, we fitted a logistic regression model with four parameters (s, 𝜶, 𝞮1, 𝞮2) to observed choices. These four parameters represent the latent transition between actions (e.g. switching from left to right) following a reversal: the latency offset s (i.e. how many trials until switching to the other side), slope α (i.e. the rate at which the switch happens), first lapse ε_1_ (before behavioral transition, immediately following a reversal), and second lapse ε_2_. (error rate after behavioral transition, once learning of reversal has occurred). **(B-D)** Fitted parameters of the choice curves during the four reversal phases. For the deterministic (100/0) reversal R1 and R2 (B-C), there was a significant sex difference in the second lapse rate parameter ε_2_, such that females had a higher error rate after the behavioral transition once they had learned the reversal, compared to male rats. During probabilistic reversal (90/10) R3 and R4 (D-E), females also tended to have higher second lapse ε_2_ compared to males. In R4 (E), females also had a sharper transition (i.e. faster switching) than males. Data points overlaid on top of the bar plots indicate mean of individual rats. Asterisks indicate significance from Wilcoxon rank sum test.

### Females initiate fewer trials than males during reversal learning

We next assessed the influences on the initiation of trials, which is often used as a proxy for attention and task engagement (17–19). For action-based reversal learning, there was a sex difference in initiation omissions (β_sex_ =-19.81, *p* = 7.91e-06) and in initiation latencies (β_sex_ =-1.17, *p* = 0.003), with females omitting more trials and taking longer to initiate trials compared to males (**Figs. 3B and 3C**). There were no sex differences in correct choice latencies (*p* = 0.77) or reward latencies (*p* = 0.88), but there was a signficant effect of sex on incorrect choice latencies (β_sex_ =-0.19, *p* = 0.04), with females taking longer than males to make an incorrect choice.

Females also omitted more trials than males (β_sex_ =-50.51, *p* = 3.27e-05) in stimulus-based reversal learning (**Fig. 3E**). There were no sex differences in any of the latency measures: initiation latencies (*p* = 0.06) (**Fig. 3F**), correct choice latencies (*p* = 0.11), incorrect choice latencies (*p* = 0.06), or reward latencies (*p* = 0.87). There was, however, an effect of reversal number on correct choice latencies (β_reversal_ =-0.04, *p* = 0.01), incorrect choice latencies (β_reversal_ =-0.05, *p* = 0.01), and reward latencies (β_reversal_ =-0.07, *p* = 0.01), with speeds increasing with each reversal.

### Use of Win-Stay and Lose-Shift is influenced by sex only in action-based reversal learning

To capture animals’ sensitivity to positive and negative feedback, we assessed their use of Win-Stay and Lose-Shift adaptive strategies during reversal learning. Males were more likely to employ a Win-Stay strategy than females (β_sex_ = 0.03, *p* = 0.02) in action-based reversal learning (**Fig. S1A**), whereas females were more likely to employ a Lose-Shift strategy than males (β_sex_ = - 0.09, *p* = 0.01; **Fig. S1B**), suggesting males were more sensitive to positive feedback, while females were more sensitive to negative feedback.

There were no sex differences in the use of adaptive strategies in either Win-stay (*p* = 0.14) or Lose-shift (*p* = 0.27) during stimulus-based reversal learning (**Fig. S1C-D**), indicating both that males and females used these strategies similarly, and that these were not the primary strategies adopted in the stimulus domain (i.e. the probabilities of Win-Stay and Lose-Shift were near chance level).

### Fitting of trial-by-trial choice data reveals sex differences in lapse rates

#### Action-based reversal learning

To capture the difference in the mechanism underlying action-based reversal learning between male and female rats, we next utilized computational models based on reinforcement learning (RL), and also examined behavioral transitions following reversals by fitting sigmoidal transition curves. For RL models, we found that the model with a passive decay parameter (*RL_decay_*) had a significantly lower AIC than the simplest model (*RL*), indicating a better fit to the choice data (*p* = 0.002, signed-rank test). Similarly, the sigmoidal curves with an additional lapse parameter (*SC2*) had significantly lower AIC compared to the simpler function (*SC1*) (*p* = 1.93e-06, signed-rank test). Comparing between the two types of models (*RL_decay_* vs. *SC2*), we found that sigmoidal transition functions had a significantly lower AIC than RL models (*p* = 2.58e-08, signed-rank test). The results were consistent when tested among male (*p* = 0.03) or female (*p* = 8.81e-05) rats separately. This suggests that the sigmoidal curve, although simpler and lacking trial-by-trial integration of reward feedback into choice probability, better described the rats’ transition behavior on each day of the reversal. Consistent with this interpretation, we found overall small values of the estimated learning rates in the RL model (*RL_decay_*), indicating slow rates of learning in both male and female rats (**Fig. S2**), especially during the first day of the reversal (M±SEM across subjects: male: 0.055±0.017; female: 0.066±0.024; **Fig. S3**). Comparing the best model (*SC2*) between male and female rats, we found that male choice behavior was overall more predictable as indicated by lower AIC (*p* = 5.25e-06, rank sum test).

Given these results, we next investigated the estimated parameter of the best-fitting sigmoidal function (SC2) (**Fig. 4A**). Overall, the lapse rates were significantly larger prior to the behavioral transition point compared to after the transition for both males (*p* = 3.56e-27) and females (*p* = 6.23e-24; mixed-effects regression on ε_1_–ε_2_), consistent with the notion that rats’ choice behavior is more noisy before transitioning to the other choice option after reversal. Yet, analysis using mixed-effects GLMs revealed that there was a significant sex difference in the second lapse rate (ε_2_), with females exhibiting a higher lapse rate after the behavioral transition point compared to males (β_sex_ = 0.05, *p* = 0.01; **Fig. 4B**). No significant main effect of sex was observed for the other parameters, after corrections for multiple comparisons (offset *s*: *p* = 0.03; slope σ: *p* = 0.08; first lapse rate ε_1_: *p* = 0.15). These results indicate that the behavioral tendency that most distinguishes males and females is the post-transition period. To confirm this possibility directly, we compared the performance of two groups after the behavioral transition point, estimated by the offset *s* (i.e., comparing *P*(*correct_Trials > s_*) within each session). We found a significant main effect of sex (β_sex_ = - 0.03, *p* = 0.01), with the female rats showing lower choice accuracy after transitioning to the better choice option. Repeating the same analysis for the number of initiation omissions (i.e., *P*(*omit_Trials>s_*)), we again found a significant sex difference (β_sex_ = 0.09, *p* = 2.62e-06). Consistently, we found that the second lapse rates were significantly correlated with the number of omissions both in females (*r* = 0.53, *p* = 2.42e-10) and in males (*r* = 0.39, *p* = 5.79e-05). This result is consistent with the interpretation that post-transition disengagement is a common state that underlies both lapse and omitted trials.

#### Stimulus-based reversal learning

We next investigated the fit to the animals’ choice data during stimulus-based task using the same RL models and sigmoidal transition curves. The results were overall consistent with the action-based task, such that the sigmoid function with two lapse parameters (*SC2*) best accounted for the choice behavior than both single-lapse sigmoid function (SC1) and the better RL model based on AIC (signed-rank test; *SC2* vs. *SC1*: *p* = 0.02; *SC2* vs. *SC2* vs. *RL_decay_*: *p* = 0.002). However, when comparing the quality of fit to the action-based task, we note that the fit to stimulus-based task was significantly worse even for the best model, offering minimal benefit over the null model with chance prediction (McFadden’s R^2^ for *SC2* in stimulus-based task: R^2^ = 0.13; **Supplementary Table 1**). For this reason, we did not follow-up with the analysis on the estimated parameters of the model for the stimulus-based task.

## Discussion

We evaluated sex differences in the context of flexible learning using stimulus-and action-based reversal learning tasks, which required animals to associate either a visual stimulus or spatial location with an outcome. Animals experienced multiple reversals that varied in probabilistic reward outcomes (first 100/0, then 90/10) and were tested on several measures, including learning and accuracy, task engagement (e.g. attention), motivation, and sensitivity to positive and negative feedback. We found consistent sex differences in rewards collected and trial initiation omissions across both stimulus-and action-based reversal learning. Using a reinforcement learning model and sigmoidal transition curves to fit choice data and characterize transitions between different behavioral states, we found females had a higher transition-specific lapse rate and poorer choice accuracy after the transition point, once there was no need for further learning.

Consistent with prior literature we did not find a sex difference in accuracy during discrimination learning (3, 13, 20, 21) irrespective of domain (i.e., actions or stimuli). All animals exhibited more difficulty learning the stimulus-based task indicated by accuracy maximally around chance level, similar to previous reports (9, 14). Only a handful of groups have conducted cross-modal comparisons of discrimination on reversal learning, finding that generally rats acquire olfactory associations faster than visual ones (22, 23). We think this learning domain asymmetry may have more to do with poor transfer of gains *across* sessions in stimulus-based learning, not a lack of trial-by-trial improvement *within* a session. Increasing the number of trials per session to 2x what we administered here is expected to greatly improve acquisition by amassing, not distributing, the learning.

Our finding that females generally omit more trials than males has been corroborated in several decision-making tasks, including the 5-Choice Serial Reaction Time Task (5-CSRTT) and the Rat Gambling Task (rGT). Previous reports-one in the 5-CSRTT (18) and another using the rGT (24) find females omit more trials than males. That we observed this pattern in both stimulus and action-based reversal learning suggests it is a generalizable phenomenon across domains. Additionally, that females collect fewer rewards and omit more trials across both domains (even though they exhibited much better learning in the action-based task) suggests that task engagement is at least partially dissociated from accuracy. To our knowledge there has only been one study that has investigated omissions in the context of flexible learning using an operant paradigm featuring both spatial and visual light-based cues, which reported that females omitted more trials and committed more perseverative errors compared to males (25).

There are likely a variety of factors that influence an animal’s ability to engage in the task (e.g., fatigue, boredom, satiety, hunger, hormones). One should consider how these internal states influence motivation and task engagement since they can also impact task performance. The role of fatigue has been mostly studied in tasks involving effort-based decision-making (26) in which animals have to exert physical effort, or in sleep studies where the animals are deliberately sleep deprived prior to being tested on decision-making paradigms (27). However, given that we did not find sex differences in either choice or reward latencies, which are reliable measures of processing speed and motivation (9, 19), fatigue and a lack of motivation are not likely explanations.

Another set of performance measures that are commonly used to assess sensitivity to positive and negative feedback are the adaptive strategies of win-stay and lose-shift, respectively. We found that males employ the win-stay strategy more than females for the action-based reversal learning task, which is consistent with prior studies that males exhibited more win-stay, and less lose-shift, compared to females using the Iowa Gambling Task (12, 28). Differences in modality or learning domain, as well as uncertainty and volatility of the environment can influence use of adaptive strategies. For example, Chen et al. (2021) found no sex differences in win-stay or lose-shift when using a visual-based restless bandit task, but found that females employed win-stay more than males during exploratory states, in a spatial-based restless bandit task. Taken together, the sex differences we found in trial omissions cannot be fully explained by sex differences in learning accuracy, satiety, or use of adaptive strategies, but does not rule out the possibility of boredom or ennui, leading to task disengagement.

In order to evaluate this further, we used computational methods based on two conventional reinforcement learning (RL) models (14) to fit the animals’ choice behavior and estimate parameters (e.g., learning rate, inverse temperature, decay rate). The fit to the choice behavior revealed an overall slow rate of reversal learning, suggesting that the shift in choice preference after reversals occurred gradually in both male and female rats. Consistent with our previous findings (14), we also found females had a lower decay rate compared to males, suggesting they retained the memory of the unchosen option’s past value for a longer time. Accordingly, we tested sigmoidal functions to characterize animals’ choice as a gradual shift in preferred options.

In particular, inspired by the previous study (13) which utilized sigmoidal transition curves to characterize post-reversal dynamics, we tested the extension of the sigmoidal curve function that accommodates distinct lapse rates prior to and following the transition in reversal. We found that this generalized model (SC2) better accounted for choice behavior in both males and females, compared to RL models. From the comparison of simulated choice behavior using the fitted parameters, the standard RL models overestimate the animals’ speed of transition and resulting performance (**Fig. S4**). However, these results alone do not necessarily rule out the presence of error-driven learning like RL, as sigmoid transition curves lack concrete mechanisms for generating choice behavior. Rather, these two types of models capture different aspects of behavior. Future modeling studies could draw on both types of models to examine more nuanced mechanisms that consider the effect of task disengagement on learning and choice. This could directly address, for example, whether choice omissions and lapse trials are generated from the same internal state. Focusing on the post-transition period, we found that females omitted more trials and exhibited poorer choice accuracy later in each reversal. Altogether, these findings suggest that females become more disengaged than males when there is not as much to learn. This disengagement may help to cope with changing reward contingencies by saving energy resources for more adaptive foraging (29) and may differ depending on the predictability and stability of the reward environment (30).

