## Supplementary Information for "Sex differences in task engagement and lapse rate during reward learning"

### Supplemental Information

| Model | - Log-likelihood (Action / Stimulus) | AIC (Action / Stimulus) | McFadden's $R^2$ (Action / Stimulus) |
|---|---|---|---|
| **SC1** | 0.2574/ 0.6417 | 0.5525 / 1.3189 | 0.629 / 0.0743 |
| **SC2** | 0.2466* / 0.6332* | 0.5434* / 1.3138* | 0.644* / 0.0865* |
| **RL** | 0.2670 / 0.6603 | 0.5590/ 1.3443 | 0.615 / 0.0474 |
| **$RL_{decay}$** | 0.2571 / 0.6449 | 0.5517 / 1.3253 | 0.629 / 0.0697 |

***Supplementary Table 1.*** Plotted are mean negative log-likelihoods, AIC, and McFadden's $R^2$ values of fitted models, separately for each task during reversal phases, R1-R4. (AO: action-based; SO: stimulus-based). Asterisk indicates significance between the best and second-best model within each task (signed-rank test, $p < .05$). Overall, the sigmoid transition function with two lapse rates (*SC2*) best accounted for the animals' choice behavior. See **Methods** for the full descriptions of each model.

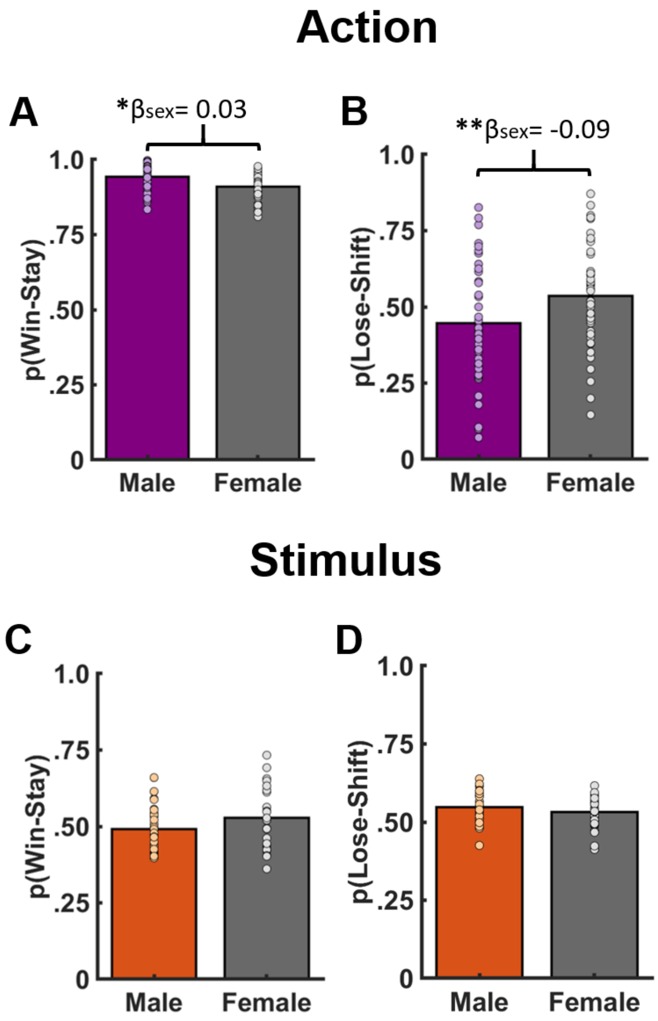


***Supplementary Figure S1. Males are more sensitive to positive feedback, females are more sensitive to negative feedback in action-based, but not stimulus-based, reversal learning.*** **(A)** Males employed the win-stay strategy more than females. **(B)** Females employed the lose-shift strategy more than males. **(C-D)** There was no significant sex difference on the use of win-stay or lose-shift strategy on any reversals for the stimulus-based task. *p<0.05, **p<.01

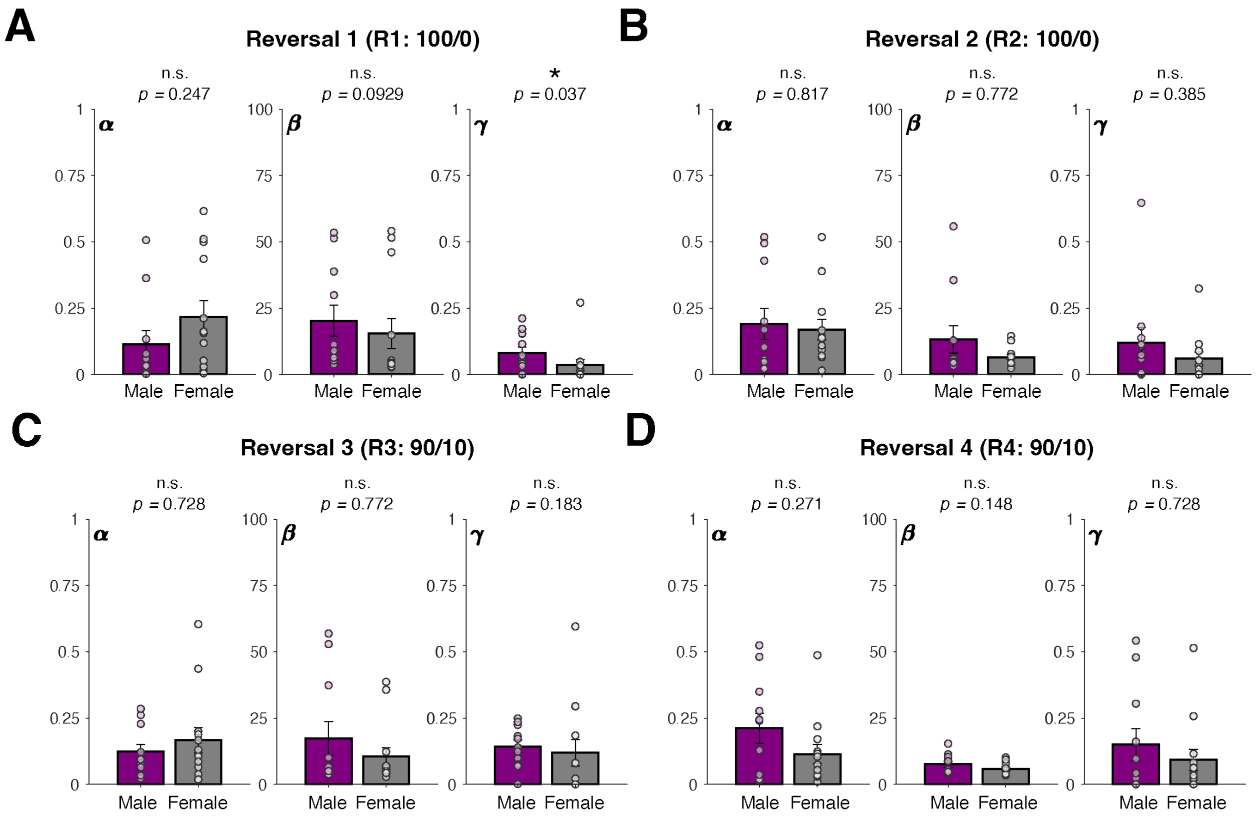


***Supplementary Figure S2. Distribution of fitted parameters in the RL model across sessions during action-based reversal learning***. Plotted are estimated parameters of the best RL model (*$RL_{decay}$*) during the four reversal phases. α: learning rate; β: inverse temperature; γ: decay rate for unchosen options. Data points overlaid on top of the bar plots indicate mean of individual rats. Asterisks indicate significant difference between male and female rats using rank-sum test. Female rats tended to have lower rate of decay on the first reversal, indicating less forgetting of unchosen values compared to male. No other significant sex differences were observed in the model parameters.

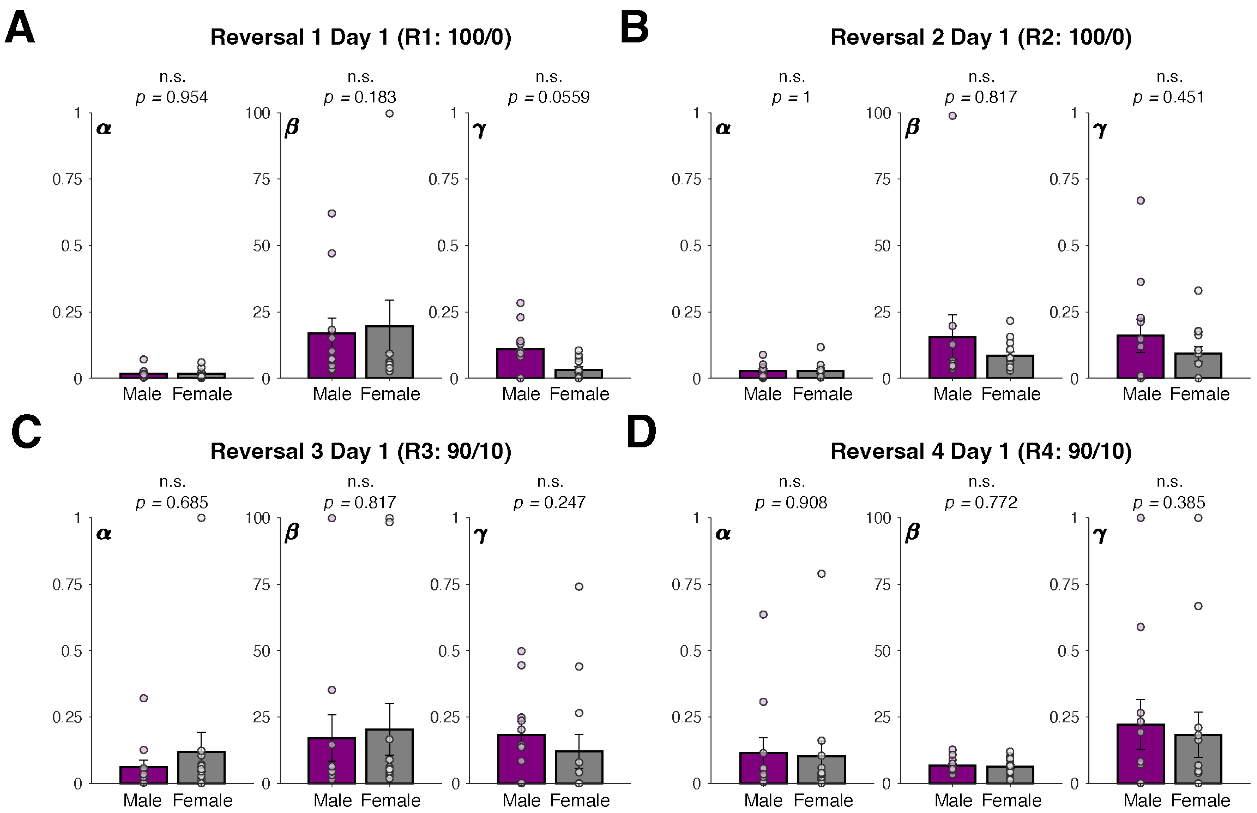


***Supplementary Figure S3. Distribution of fitted parameters in the RL model during the first days of the reversals during action-based reversal learning***. Plotted are estimated parameters of the best RL model ($RL_{decay}$) during the four reversal phases, shown for only the first day of each phase. Convention is the same as in **Fig. S2**.

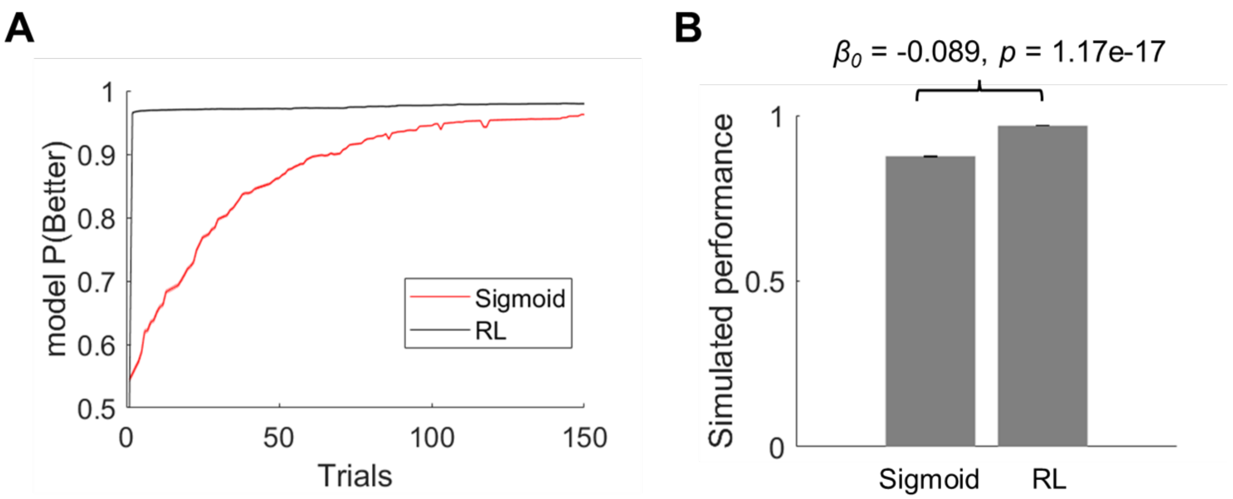


***Supplementary Figure S4. Comparison of simulated choice behavior in the Action-based task using sigmoidal and RL models.*** **(A)** Plotted are averaged model-generated choice probabilities for the better option across simulated sessions (100/0 reward schedule), shown for the first 150 trials. The best respective model for sigmoid and RL were simulated (*SC2* and $RL_{decay}$). Each deterministic session was simulated 100 times with the estimated parameters of each model. The RL model required only a few trials to reverse their choice option, overestimating the transition speed. **(B)** Comparison of simulated performance from two models. Sigmoid model had significantly lower performance (mixed-effects regression on paired difference in performance, $(P_{sigmoid} - P_{RL}) \sim 1 + (1|SessionID)$; *reported is the intercept* $\beta_0$ and its p-value). With faster transition to the better option, RL model overestimates performance.
